# Closed-loop sacral neuromodulation for bladder function using dorsal root ganglia sensory feedback in an anesthetized feline model

**DOI:** 10.1101/2020.05.02.074484

**Authors:** Zhonghua Ouyang, Nikolas Barrera, Zachariah J. Sperry, Elizabeth C. Bottorff, Katie C. Bittner, Lance Zirpel, Tim M. Bruns

## Abstract

Overactive bladder patients suffer from a frequent, uncontrollable urge to urinate, which can lead to a poor quality of life. We aim to improve open-loop sacral neuromodulation therapy by developing a conditional stimulation paradigm using neural recordings from dorsal root ganglia (DRG) as sensory feedback. Experiments were performed in 5 anesthetized felines. We implemented a Kalman filter-based algorithm to estimate the bladder pressure in real-time using sacral-level DRG neural recordings and initiated sacral root electrical stimulation when the algorithm detected an increase in bladder pressure. Closed-loop neuromodulation was performed during continuous cystometry and compared to bladder fills with continuous and no stimulation. Overall, closed-loop stimulation increased bladder capacity by 13.8% over no stimulation (p < 0.001) and reduced stimulation time versus continuous stimulation by 57.7%. High-confidence bladder single units had a reduced sensitivity during stimulation, with lower linear trendline fits and higher pressure thresholds for firing observed during stimulation trials. This study demonstrates the utility of decoding bladder pressure from neural activity for closed-loop control of sacral neuromodulation. An underlying mechanism for sacral neuromodulation may be a reduction in bladder sensory neuron activity during stimulation. Real-time validation during behavioral studies is necessary prior to clinical translation of closed-loop sacral neuromodulation.

## 1. Introduction

Overactive bladder (OAB) is a dysfunction that affects millions of people worldwide. Patients suffer from frequent urinary urgency, with or without incontinence [1], leading to a variety of side effects such as poor sleep, declined mental health, and a low quality of life [2]. Conservative therapies such as anticholinergic drugs and intravesicular Botox injections are both associated with undesirable side effects, leading to low patient compliance [3], [4], and anticholinergics are also associated with an increased risk of dementia [5]. Currently there is no pharmaceutical therapy that permanently reduces or eliminates the symptoms without side effects [2].

Sacral neuromodulation (SNM) is a standard clinical treatment for OAB after conservative approaches such as behavioral modification and pharmaceuticals fail [6]. A recent study reported that 82% of patients discontinued OAB medication after SNM treatment for at least 22 months [7]. In SNM, a stimulation lead is placed near the S3 or S4 sacral nerve in a minimally invasive surgery. SNM is applied constantly at 14 Hz to reduce the symptoms of OAB [8]. SNM generally has high success rates, defined as the percentage of patients with at least a 50% improvement, of 61-90% for improving overactive bladder symptoms [9]. SNM consistently has less frequent side effects than Botox for refractory OAB [10], [11], but studies have reported differences in which treatment yields higher success rates [10]–[12]. Significantly higher success rates have been reported for SNM as compared to standard medical treatment [13], however there is a lack of long-term comparisons between SNM and pharmacological therapies.

SNM patients can experience relapse of symptoms or loss of efficacy over time [14]. Continuous stimulation can facilitate habituation of neural pathways over time [15], [16], which may contribute to the relapse of symptoms. Pre-clinical studies have demonstrated that conditional or closed-loop stimulation of peripheral nerves may offer greater clinical benefit by modulating bladder function only when necessary, leading to increased bladder capacity [17], [18]. Similarly, pilot studies with neurogenic and non-neurogenic patients have shown the potential for conditional dorsal genital nerve stimulation to improve bladder capacity over continuous stimulation [19]–[21]. Studies with non-continuous SNM, using various fixed duty cycles, have reported similar positive outcomes for OAB and incontinence patients as continuous stimulation [22]–[26]. Patients in these studies usually report a preference for lower duty cycle, non-continuous stimulation [22], [25], [26]. In a study in which patients controlled conditional SNM, 10 of 16 participants maintained bladder improvements, though this approach requires patient intervention. A reliable method for sensing bladder events that does not require patient intervention is needed [27], [28]. A sensing method that is adjacent to a stimulation site may be particularly beneficial.

Bladder sensory signals can be observed at sacral dorsal root ganglia (DRG) [29]. In addition to physical proximity to the sacral neuromodulation location, sacral-level DRG contain afferent-only signals from the detrusor muscle and urethra, via proximal pelvic and pudendal nerve fibers [30]. In this study, we used sacral-level DRG as a recording site to estimate bladder pressure and the onset of bladder contractions in real-time [31] to automatically trigger closed-loop neuromodulation in anesthetized, healthy cats. This is the first study to examine the effect of closed-loop SNM on bladder capacity using neural signals as feedback. While from a pre-clinical research perspective, bladder capacity is usually considered as the most important performance metric, in this study we also evaluate non-voiding bladder contractions. Sensory neurons activated during bladder pressure increases and voiding contractions are also usually activated during non-voiding contractions [31]. These non-voiding contractions may contribute sensations of urgency in OAB, and are therefore undesirable. We hypothesize that closed-loop stimulation increases bladder capacity over no-stimulation trials and reduces the frequency of non-voiding contractions (NVCs) to the same extent as continuous stimulation while applying significantly less stimulation. Although this study was performed in an anesthetized, non-OAB animal model, it shows the potential for closed-loop SNM to improve bladder function that should be explored further with a long-term study.

## 2. Methods

### 2.1 Animals

All procedures were approved by the University of Michigan Institutional Animal Care and Use Committee (IACUC), in accordance with the National Institute of Health’s guidelines for the care and use of laboratory animals. Five adult, domestic, short-hair male cats (0.99 ± 0.27 years old, 4.70 ± 0.57 kg, Marshall BioResources, North Rose, NY) were used in this study (designated as experiments 1–5). Cats were used due to their high relevance to human physiology and their long history of study in bladder neurophysiology [32], [33]. Prior to use, animals were free-range housed with up to 3 other cats in a 38 m^2^ room with controlled temperature (19-21 °C) and relative humidity (35-60%), food and water available ad lib, and a 12-hour light/dark cycle. Animals received enrichment via staff interaction and toys.

### 2.2 Surgical procedure

As in prior studies [31], [34], animals were anesthetized with a mixture of ketamine (6.6 mg/kg), butorphanol (0.66 mg/kg), and dexmedetomidine (0.011 mg/kg) administered intramuscularly. Animals were intubated and subsequently maintained on isoflurane anesthesia (0.5-4%) during surgical procedures. Respiratory rate, heart rate, end-tidal CO_2_ , O_2_ perfusion, temperature, and intra-arterial blood pressure were monitored continuously using a SurgiVet vitals monitor (Smiths Medical, Dublin, OH). Fluids (1:1 ratio of lactated Ringers solution and 5% dextrose) were infused intravenously via the cephalic vein at a rate of 10 ml/hr during surgery as needed. A 3.5 Fr dual-lumen catheter was inserted to the bladder through the urethra for fluid infusion and pressure monitoring. The urethra was not ligated. A midline dorsal incision was made to expose the L7 to S3 vertebrae and a partial laminectomy was performed to access sacral DRG. A lab-fabricated bipolar stimulation cuff (1.5 or 2 mm inner diameter) was placed on the left or right S1 root encompassing both the sensory and motor branches.

Two iridium oxide microelectrode arrays for neural recordings (4×8 configuration; 1.0 mm shank length; 0.4 mm shank pitch; Blackrock Microsystems, Salt Lake City, UT) were implanted into (1) left and right S1 DRG or (2) S1 and S2 DRG on the same side using a pneumatic inserter (Blackrock Microsystems). Array reference wires were placed near the spinal cord and ground wires were attached to a stainless steel needle inserted below the skin (lateral and caudal to the laminectomy incision site). At the conclusion of surgical procedures, prior to experimental testing, animals were transitioned to intravenous alpha-chloralose (C0128, Sigma Aldrich; 70 mg/kg induction; 20 mg/kg maintenance). This transition was at least six hours after induction, and we expect that there were no residual effects on bladder function due to the induction dosing of ketamine, butorphanol, or dexmedetomidine. Analgesia was augmented with 0.01 mg/kg buprenorphine every 8–12 hours subcutaneously.

### 2.3 Closed-loop SNM system

Prior to the main cystometry experiments, stimulation parameter optimization was performed in preliminary experiments. In isovolumetric trials, 5 Hz (200 µs biphasic pulse width) stimulation on the S1 root was more effective at inhibiting bladder non-voiding and voiding contractions at 2 times the motor threshold (MT) for scrotum, anus or tail twitching, compared to 2, 7, 10, 15, and 33 Hz. 5Hz was thus selected for all experiments in this study, as has been performed in prior cat SNM studies [35]–[37], though it differs from the standard 14 Hz for clinical SNM [8], [13]. The stimulation amplitude used for cystometry trials was varied across experiments, within 1-4 x MT (0.10 mA-0.72 mA), depending both on how effective bladder contractions were suppressed and the extent of movement artifacts that were caused by stimulation. Experiments 1 and 5 used 1-1.5 x MT, experiments 2 and 3 used 4 x MT, and experiment 4 used 2 x MT. It has been shown that supra-MT stimulation in anesthetized animals is necessary for effective study of SNM [37], [38], in contrast to clinical SNM which is typically set based on the sensory threshold [22], [39] which may vary with respect to MT.

Sacral nerve stimulation and microelectrode array recordings were integrated through the Medtronic Summit Research Development Kit (RDK) to deliver closed-loop stimulation. The RDK is comprised of an Olympus RC+S (B35300R) Implantable Neural Stimulator (INS) connected to a four-electrode stimulation lead (Medtronic Model 3889), the Summit Application Programming Interface (API), and other supporting hardware components including a Research Lab Programmer (RLP, a tablet mainly for setting stimulation parameters and safety limits), Clinician Telemetry Modules (CTMs) that enable wireless connection between the INS and the research host computer (for delivering closed-loop stimulation) or the RLP, a Patient Therapy Manager (PTM, for charging the INS and parameter setting), and a Recharge Therapy Manager (RTM) that enables inductive charging of the INS through the PTM. This full system was not specifically necessary to perform closed-loop stimulation during these anesthetized experiments, as we have demonstrated in a previous pilot experiment [31]. One of our goals for this study was to demonstrate feasibility with a wireless human use system [40] as a steppingstone for behaving preclinical experiments with a fully implantable system towards clinical translation.

The two stimulation cuff lead wires were attached to two of the four electrodes on the INS lead with modelling clay, which served both as an adhesive and an insulator for the connection. A setup diagram is shown in Figure 1. The Summit RDK software package allows programmatic control of the INS through a MATLAB interface, in which a previously developed Kalman filter algorithm [31], [34] was implemented to decode bladder pressure (as a control signal) from the neural recordings (collected at 30 kHz per channel through the Ripple Grapevine system and Trellis software). The neural recordings were processed into threshold crossings with a positive and negative dual threshold at ±3.5-5.5 × root-mean square of the raw signal. A spike only needed to cross one of the thresholds to be captured. The algorithm extracted unsorted threshold crossing firing rates using a non-sliding 1-second interval window from recording channels and combined them with a state-dependent model to estimate the bladder pressure using a weighted average method. DRG microelectrode channels with a firing rate to bladder pressure correlation of greater than 0.7 were included in the model. If no more than one channel had a correlation greater than 0.7, then this cutoff was reduced in 0.1 increments until there was at least two channels used by the model. A cross-channel invalidation method was applied before the firing rates were calculated to remove threshold crossings that simultaneously appeared on over 90% of the channels, to minimize the effect of stimulation artifacts.

**Fig 1.**
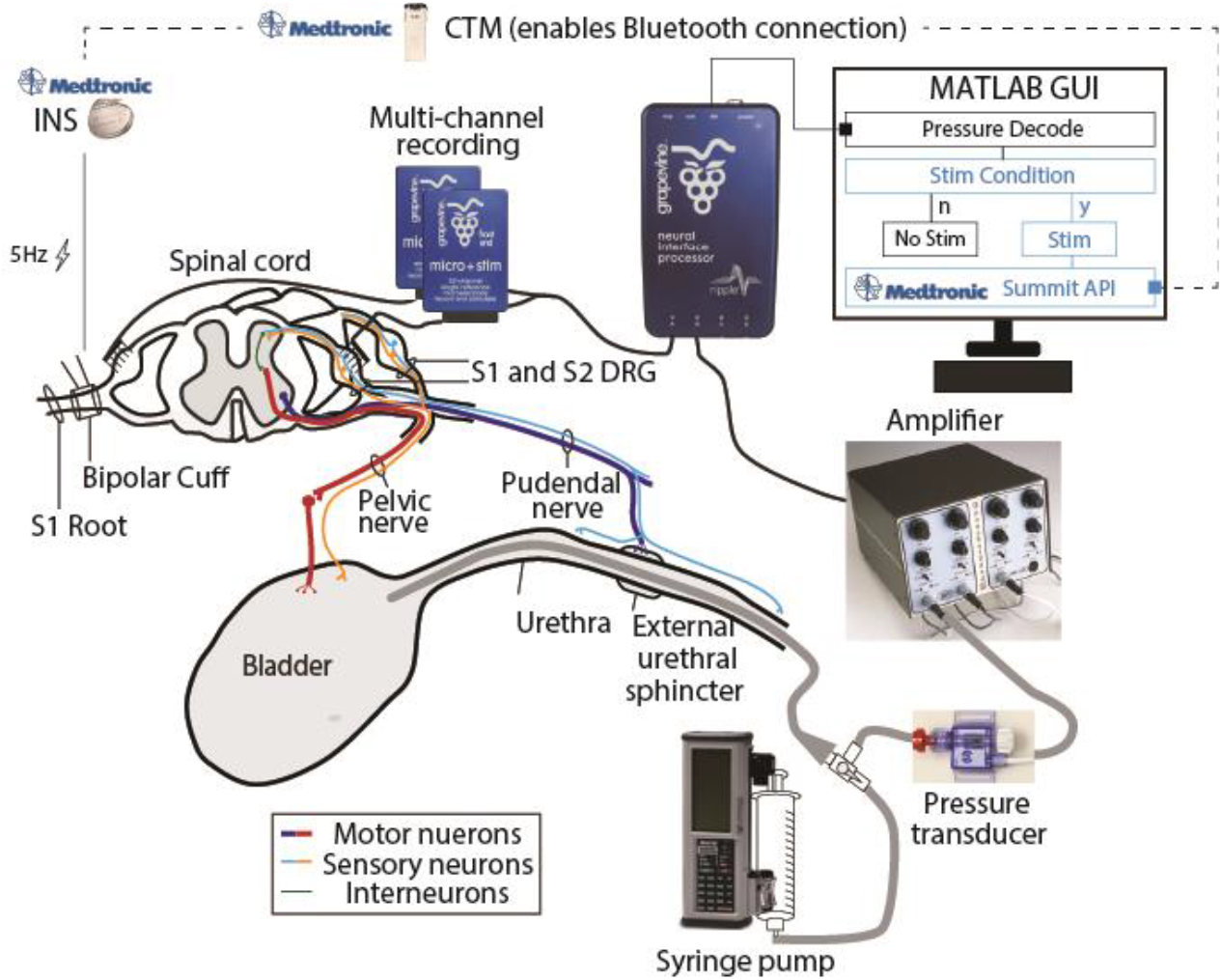
Illustration of the testing setup. DRG neural recordings were acquired with a Ripple Grapevine system and accompanying Ripple Trellis software via microelectrode arrays implanted in S1 and S2 DRG. Trials consisted of recording neural data and bladder pressure (monitored with a pressure transducer and amplifier) during saline infusions at a controlled rate via an intraurethral bladder catheter. Pressure data was recorded with the Grapevine system after amplification. Real-time decoding was performed in a MATLAB GUI that contains the Summit Application Programming Interface (API) that enables Bluetooth control of the Implantable Neural Stimulator (INS) through a Clinician Telemetry Module (CTM)

### 2.4 Experimental trials

Cystometry trials were performed in which the bladder was infused with 0.9% saline (warmed to 41 °C) through a 3.5 Fr urethra catheter at 2 ml/min from an empty volume to when the first leak around the urethra catheter was observed. In each trial, one of three stimulation paradigms was used: continuous (“continuous SNM”), closed-loop stimulation, or no stimulation (except in experiment 5, in which only no-stimulation and closed-loop stimulation were performed due to unanticipated experimental circumstances that limited the testing time). In closed-loop trials, stimulation was initiated based on a bladder contraction detection algorithm, indicating when the DRG decoding algorithm showed an increase in bladder pressure within a certain time window. The first two experiments were exploratory, in which the contraction detection algorithm was varied and closed-loop sacral nerve stimulation was conditionally turned on for a fixed duration of 15-60 s when an estimated bladder pressure increase of 3-10 cmH_2_O was observed from the start of a 3-6 s window. Observations from cystometry trials and additional isovolumetric trials in these two experiments and a third, separate experiment without cystometry trials were used to select the final contraction detection algorithm. In the last three experiments, the contraction detection algorithm was fixed, and closed-loop stimulation was turned on for 15 s after a 6 cmH_2_O increase in estimated pressure was observed from the start of a 4-second moving window. The primary goal of this study was to demonstrate closed-loop stimulation, we prioritized running as many closed-loop and no-stimulation trials as possible within a limited experimental time; therefore, the order of the trials was not completely random.

After each trial ended, the bladder capacity was measured as the amount of fluid in the bladder when the first leak was observed. This was done by adding the fluid volume manually emptied from the bladder (through a urethral catheter) and any leak volume collected by a weigh boat. The bladder was allowed to rest for at least 15 minutes before initiating the next cystometry trial.

### 2.5 Euthanasia

After completion of all testing, animals were euthanized with a dose of intravenous sodium pentobarbital while under deep isoflurane anesthesia, and death was ensured with a secondary method of euthanasia as approved by the IACUC.

### 2.6 Data analysis

Bladder capacity was measured in each trial and normalized to the control (no-stimulation) group average in each experiment (animal). A rank-based non-parametric Kruskal-Wallis test was used to evaluate statistical significance in bladder capacity among no-stimulation, closed-loop stimulation, and continuous stimulation across experiments and across all trials. A significance level of 0.05 was used. Post hoc pair-wise comparisons were performed with a Kruskal-Dunn test, with Bonferroni correction on p-values.

We defined bladder contractions (NVCs or voiding contractions) as bladder pressure increases of at least 6 cm H_2_O in a 4-second interval (independent from the contraction detection algorithm). The number of bladder contractions were counted for each trial and normalized to the average of the no-stimulation trials for each experiment. The timing of each bladder contraction was matched with the stimulation initiation timing, and the true positive rate was calculated by dividing the number of true positives (a bladder contraction successfully identified by the algorithm) by the total number of true bladder contractions. The average interval between bladder contractions for each trial was normalized to the no-stimulation group average for each experiment (animal). Similarly, the peak pressure (maximum pressure during voiding) for each trial was also normalized to the no-stimulation group average for each experiment (animal).

The bladder pressure decoding performance was determined using the normalized root mean squared error (NRMSE) and the correlation coefficient (R) between the measured pressure and estimated pressure [31], [34].

While reviewing data after completion of experiments, we observed that stimulation may have had an effect on neural signalling. Thus we performed an additional post hoc analysis in which we identified putative bladder single units that appeared in at least one no stimulation trial and one continuous SNM trial. The units were isolated in Offline Sorter (Plexon, Dallas, TX) with automated clustering via principal component analysis followed by manual review of snippet waveform shapes by an experienced spike sorter. Only high-confidence single units that had a clearly identifiable waveform shape were included in this analysis. For SNM trials, 5 Hz stimulation artifacts were isolated from neural activity based on clearly differentiable waveforms appearing at a fixed frequency. We confirmed that stimulation artifacts had minimal to no concealment of bladder unit snippets by reviewing artifact waveforms. To quantify the relationship between identified bladder units and pressure, the correlation coefficient, linear regression slope, and minimum pressure at which a unit started firing (pressure threshold) were determined for each unit. We determined that a unit had started to fire when a single spike was detected. The average change in these parameters from no stimulation trials to SNM trials was calculated. A limited statistical analysis was performed due to a small sample size, as described below.

## 3. Results

Overall, we performed 30 no-stimulation, 23 closed-loop stimulation, and 9 continuous SNM trials across five experiments. Bladder pressure decoding was performed in all closed-loop trials, and some of the no-stimulation and continuous SNM trials. Example testing trials in one experiment for all three conditions are shown in Figure 2.

**Fig 2.**
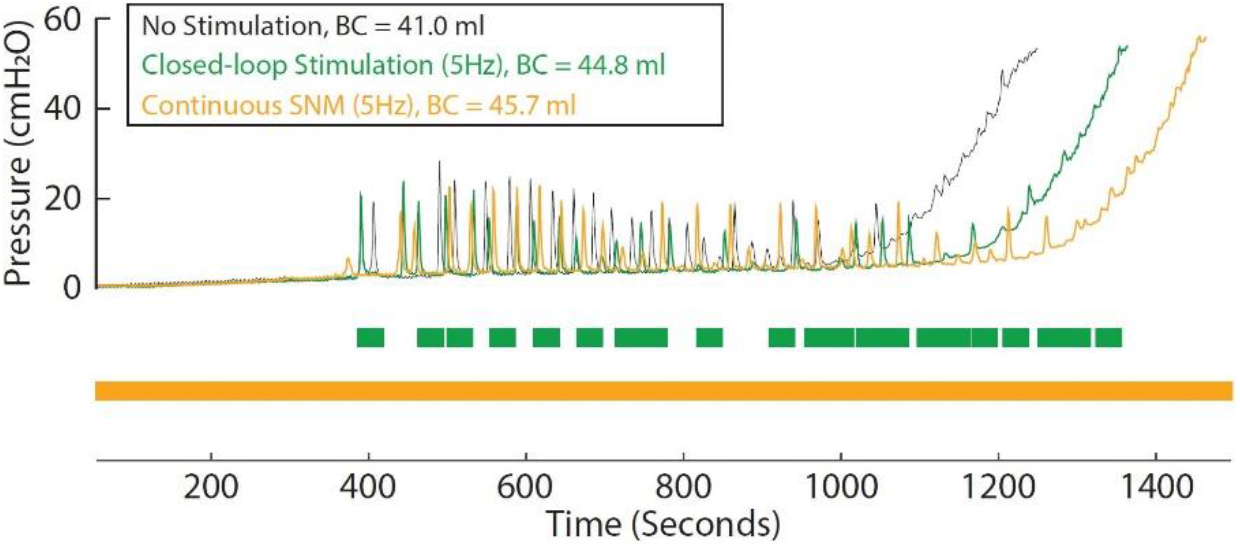
Cystometry curves for example no-stimulation, closed-loop stimulation, and continuous SNM in Experiment 2. Closed-loop stimulation and continuous SNM increased bladder capacity (BC) compared to no-stimulation trials in these examples

### 3.1 Normalized bladder capacity

To assess the effectiveness of each SNM paradigm, we calculated the ratio between the bladder capacity for each trial and the average no-stimulation capacity in an experiment across experiments and all trials for each stimulation type. We observed a 12.8 ± 4.6% mean per-experiment increase in normalized bladder capacity in the closed-loop stimulation group across all 5 experiments, and a 12.9 ± 6.5% per-experiment increase in normalized bladder capacity in the continuous SNM group in 4 experiments (no continuous SNM in experiment 5). The mean per-experiment bladder capacities were significantly different from control bladder capacities (p = 0.02 for each, Figure 3a). Across all individual trials performed, the increase in normalized capacity was 13.8% (p < 0.001, range: -12 to 34%) for closed-loop stimulation (n = 23 trials) compared to control trials (n = 30). No statistical significance was found between all continuous SNM trials (9.1%, range: -27% to 42%, n = 9, Figure 3a) and control trials (p = 0.26). Due to time limitations in each anesthetized experiment, different counts of stimulation trial types were performed in each experiment, with an emphasis placed on performing as many closed-loop and no-stimulation (for control and buffering between stimulation trials) trials as possible. Across experiments, stimulation amplitude and the animal itself did not affect bladder capacity.

**Fig 3.**
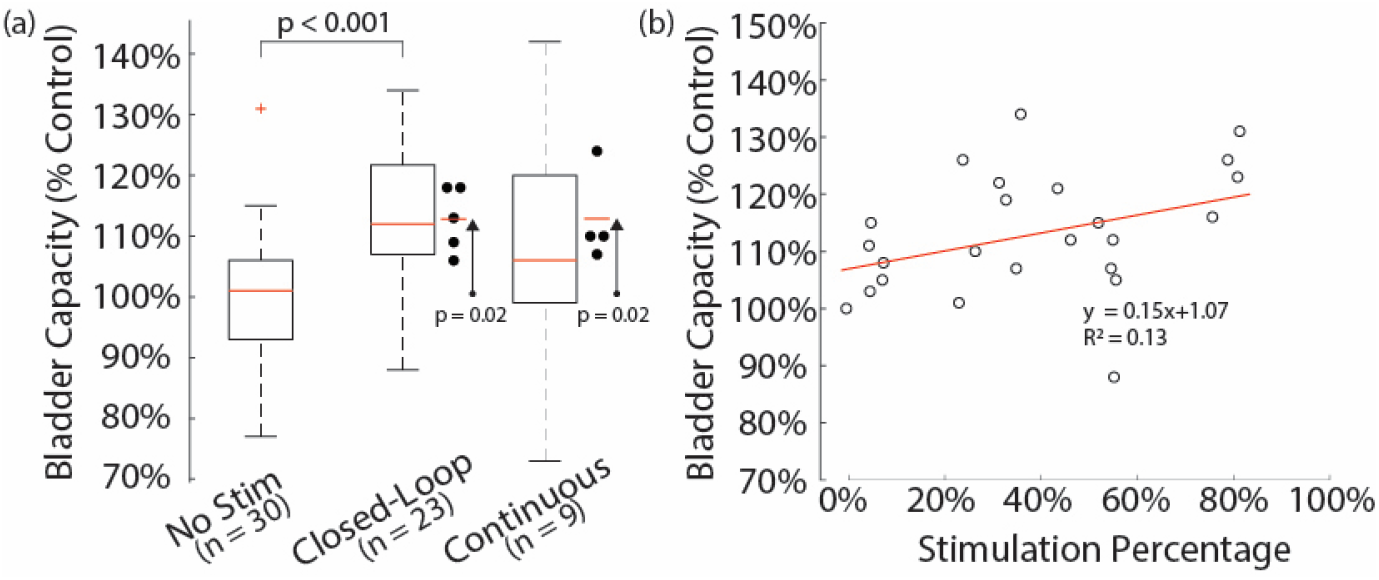
(a) Bladder capacity for each stimulation type for all trials. Circles next to Closed-Loop and Continuous boxplots give per-experiment mean values. (b) Bladder capacity against stimulation percentage for each trial. The red line is a linear fit (p = 0.15)

Across all trials, we observed a non-significant positive correlation (R^2^ = 0.13, p = 0.15) between normalized bladder capacity and stimulation percentage (total time when stimulation was on divided by total trial time, Figure 3b). While this indicates that more stimulation was associated with a stronger bladder inhibition effect, there was no difference between partial (closed-loop) stimulation and continuous SNM in terms of bladder capacity (p = 0.80).

### 3.2 Closed-loop algorithm performance

To determine bladder contraction detection accuracy, we evaluated the ratio between correctly identified contractions and total number of contractions for each trial and across each experiment. On average, 39.5 ± 12.5% (across n = 5 experiments) of the non-voiding contractions were correctly identified by the decoding algorithm, triggering stimulation. The true positive rate was 35.9% (11.4% and 48.4% for the first and second halves of the cystometry trials, averaged across n = 5 experiments). Of the stimulation bouts triggered in all trials, 51% of the stimulation occurred in the first 75% of the cystometry, while 49% of the stimulation occurred in the last 25% of the cystometry (n = 5 experiments). Figure 4 shows the distribution of stimulation by time-normalized quartiles across all 23 closed-loop trials.

**Fig 4.**
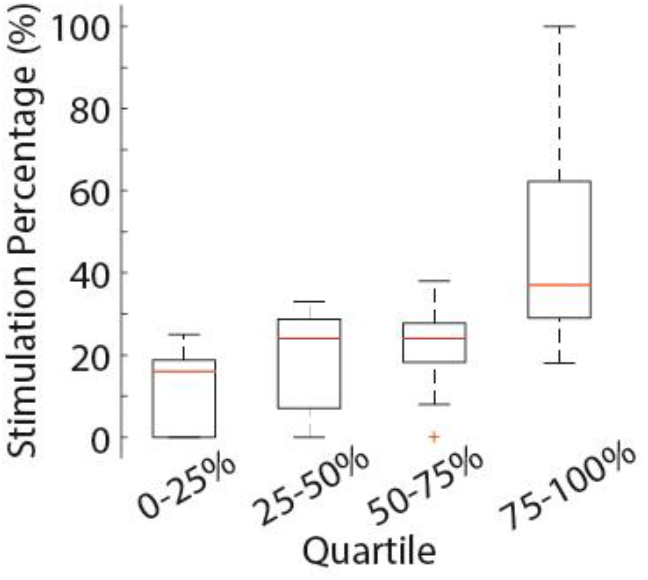
Box plots showing the percentage of overall stimulation time per closed-loop trial distributed across time-normalized quartiles

To examine the effect of stimulation on non-voiding contractions, we calculated inter-contraction intervals and contraction counts in each experimental stimulation group. Overall, closed-loop stimulation led to a 112.4% increase in the normalized non-voiding contraction interval (n = 5 experiments), while resulting in a small 3.2% decrease in the normalized number of contractions (n = 5 experiments) per trial. We observed that in some cases the start of stimulation corresponded with an NVC occurrence. This effect was not quantified but may have contributed to a minimal change in NVC count in the closed-loop stimulation group. While continuous SNM increased the non-voiding contraction interval by only 26.8%, it decreased the number of non-voiding contractions per void by 51.2%. The peak bladder pressure was slightly increased by closed-loop and continuous stimulation (1.5% and 3.9%). These non-voiding contraction and pressure trends were not statistically significant.

### 3.3 Decoding performance

To evaluate the decoding performance of the algorithm during stimulation, we calculated the NRMSE and R between the estimated bladder pressure and measured bladder pressure for each trial. Bladder pressure decoding was performed in real-time in each type of trial (Table 1). On average, decoding was performed with 5 DRG microelectrode channels (range: 2-11) in each experiment. As predicted, closed-loop stimulation trials had an increase in NRMSE and a decrease in R for bladder pressure estimation, as additional channel threshold crossings were detected during stimulation. While a cross-channel invalidation method was applied to remove threshold crossings that appeared on over 90% of the channels at the same time, we still observed an overestimation of the bladder pressure during stimulation. This was mostly due to stimulation-driven units and stimulation artifacts that appeared on fewer than 90% of the channels. In addition, a positive, non-significant trend was observed between normalized bladder capacity and the decoded pressure correlation coefficient R (Figure 5).

**Table 1.** Decoding performance by NRMSE and R across stimulation trials

|  | NRMSE |  |  | R |  |  |
| --- | --- | --- | --- | --- | --- | --- |
|  | No Stim | CLS | Continuous SNM | No Stim | CLS | Continuous SNM |
| Mean | 0.19 | 0.29 | 0.18 | 0.83 | 0.62 | 0.78 |
| St. Dev. | 0.07 | 0.17 | 0.04 | 0.18 | 0.22 | 0.01 |
| n | 9 | 23 | 3 | 9 | 23 | 3 |
| % Increase |  | 55.27% | -4.57% |  | -25.91% | -7.07% |

(CLS = Closed-loop Stimulation)

**Fig 5.**
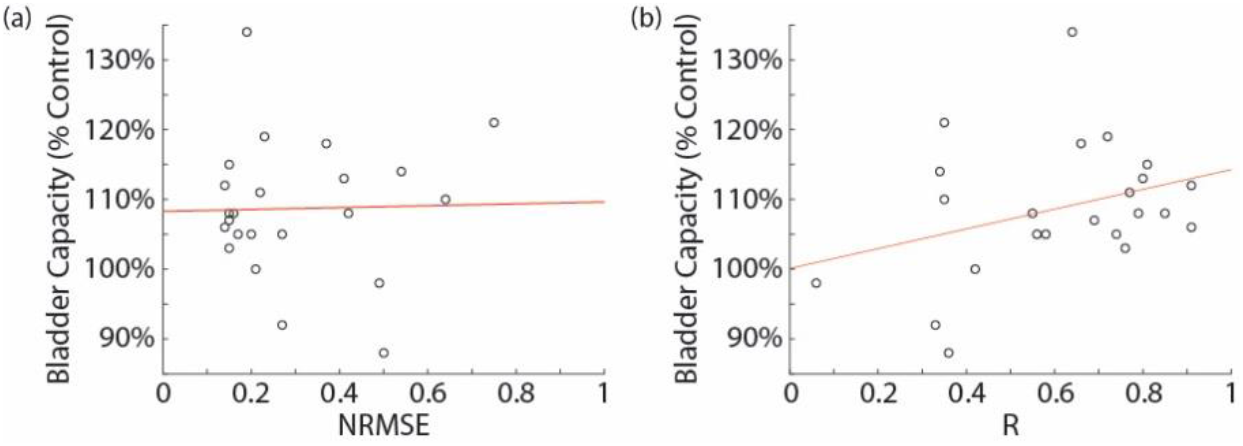
(a) Normalized bladder capacity vs. NRMSE (R^2^ = 0.02, p = 0.52) (b) Normalized bladder capacity vs. R. (R^2^ = 0.08, p = 0.18)

### 3.4 Single unit analysis

To investigate a potential interaction between SNM and bladder sensory neurons, we examined the relationship between their neural signals and bladder pressure during stimulation and no-stimulation conditions. In experiments 1-4, eight bladder units that appeared in at least one no-stimulation and one continuous SNM trial were identified with manual spike sorting. While overall we observed a much larger number of DRG microelectrode channels with bladder activity, only single units that were clearly distinguishable from stimulation artifacts were included in this analysis. On average, the correlation coefficient between the firing rate of these units and the bladder pressure during continuous SNM trials was 9.1 ± 57.2% lower than during no-stimulation bladder fills (p = 0.84, Wilcoxon Signed Rank Test). The slope of linear regression trendlines between the single unit firing rate and bladder pressure in SNM trials was 35.5 ± 47.1% lower than no-stimulation trials (p = 0.11). The minimum pressure at which bladder units first fired in SNM trials was 4.7 ± 5.5 times higher than in no-stimulation trials (p = 0.04). For all bladder unit trials pooled together by stimulation type, continuous stimulation trials had a significantly higher pressure threshold (3.9 ± 3.7 cm H_2_O, range 0.8-10.2 cm H_2_O, median 2.3 cm H_2_O, p = 0.024) than no-stimulation trials (1.8 ± 2.4 cm H_2_O, range 0.1-8.7 cm H_2_O, median 0.9 cm H_2_O) according to a Mann Whitney U test. Two of these bladder units are shown in Figure 6, with a representative 3-second interval showing differentiation of bladder units and SNM artifacts in Figure 6c. The parameters for each bladder unit are presented in Table 2.

**Table 2.** Bladder unit change in correlation coefficient, linear regression slope, and pressure threshold change with stimulation.

| Exp | Unit # | R Change | Slope Change | Pressure Threshold Change | No-Stim Trials | SNM Trials | Stimulation & recording electrode relative spinal nerve locations |
| --- | --- | --- | --- | --- | --- | --- | --- |
| 1 | 1 | -83.8% | -97.6% | 389.2% | 3 | 1 | Opposite |
| 1 | 2 | 3.4% | -43.8% | 37.7% | 3 | 1 | Opposite |
| 1 | 3 | -1.2% | -16.8% | 85.0% | 3 | 1 | Opposite |
| 1 | 4 | 4.1% | -1.3% | 65.2% | 2 | 1 | Opposite |
| 1 | 5 | 101.8% | 46.9% | -80.0% | 3 | 1 | Opposite |
| 2 | 1 | -74.5% | -92.6% | 1337.0% | 3 | 1 | Opposite |
| 3 | 1 | -23.6% | -42.0% | 1136.4% | 4 | 3 | Opposite |
| 4 | 1 | 1.1% | -37.1% | 764.6% | 1 | 1 | Same |
|  | Mean | -9.1% | -35.5% | 466.9% |  |  |  |
|  | St. Dev. | 57.2% | 47.1% | 546.5% |  |  |  |

**Fig 6.**
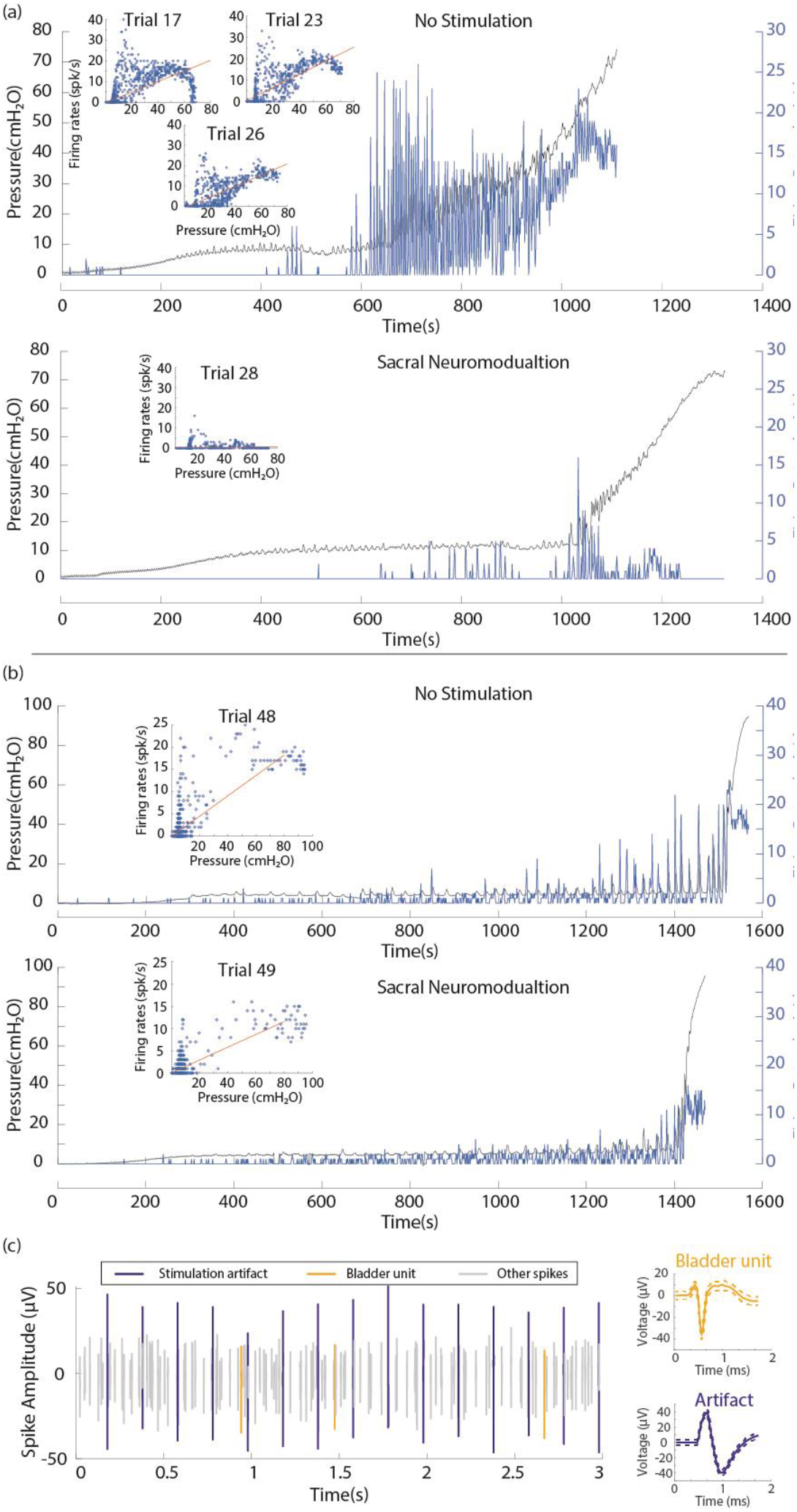
Two examples of sorted bladder single units, from experiments 1 (a) [unit 1] and 4 (b), which demonstrate a reduction in sensitivity to bladder pressure changes during continuous SNM. Inset figures plot firing rate against pressure at each calculation interval, with linear regression trend lines overlaid in red. For (a), the no-stimulation Trial 26 is plotted against time. (c) Left: Raster plot of sorted threshold crossings showing a bladder unit, stimulation artifacts, and other crossings during an example SNM trial [exp. 1, unit 2], demonstrating differentiation of signals. Right: averaged bladder unit waveform (yellow) and stimulation artifact waveform (blue)

## 4. Discussion

In this study, we explored the short-term efficacy of closed-loop sacral nerve stimulation for increasing bladder capacity in an anesthetized animal model. If translatable, our results suggest that closed-loop SNM could have a potential clinical impact by providing automated, individualized therapy that is linked to objective, physiological signals and a reduction in battery drain due to stimulation. While a reduction in battery use is primarily advantageous for primary cell devices, in the context of rechargeable devices that remain implanted in the patient for longer periods of time [41], closed-loop SNM may increase the recharge interval and improve chronic maintenance of therapy. Our study demonstrates that closed-loop stimulation may allow for this by providing the same or improved performance while applying stimulation at a fraction of the time. A reduction in total stimulation time may reduce nerve habituation over time and preserve the responsiveness of the stimulation target. However, the control system necessary to record neural signals and implement closed-loop control will require battery power. Systems for low-power neural recording and closed-loop control are being developed for other neurotechnology applications [42]–[44], and could be implemented for bladder control as the technology matures. Neural recording and bladder state estimation could be turned off for a period of time after voiding events, based on an individual’s typical voiding interval and recent activity, to further save battery power. Additionally, our results suggest that SNM desensitizes bladder sensory neurons to changes in bladder pressure. This potential mechanism of SNM may be an important contributor to its therapeutic benefit and warrants further exploration.

We successfully achieved closed-loop SNM by integrating real-time bladder pressure decoding from DRG with the Medtronic Summit RC+S stimulation system, which has been used in clinical research [40]. We demonstrated this full integration with in-vivo experiments and showed that closed-loop stimulation had at least the same level of effectiveness as continuous SNM in increasing bladder capacity (Figure 3a), however stimulation was only applied 42.3% of the time, on average. The average normalized increase in bladder capacity for closed-loop across experiments (12.8%) and all trials (13.8%) was not significantly different than continuous SNM across experiments (12.9%) and all trials (9.1%) in this study. These per-experiment and per-trial mean increases in bladder capacity were similar to a previous study (13.4%) that also performed continuous sacral root stimulation with a chloralose-anesthetized feline model [45]. Zhang et al found that stimulating the dorsal side of the subdural sacral root increased bladder capacity in cats by 64% but used a different stimulation frequency (10Hz) [37]. Their experimental model also had a more invasive procedure than ours, in which one ureter was cut and tied, while the other was used for draining. Jezernik et al. observed that stimulating the dorsal root eliminated reflex bladder contractions [46]. Stimulating only the dorsal side is challenging in humans considering its proximity to the spinal cord and tight space in the spinal column. It may be possible to target the dorsal root or DRG for stimulation, instead of the whole spinal root as for SNM, using the percutaneous lead insertion method developed for DRG stimulation for chronic pain [47]. However, the standard SNM surgical procedure, in which the lead is inserted directly through the S3 foramen to target the spinal nerve, is a simpler procedure.

Closed-loop studies have shown that stimulation on peripheral nerves during voiding, non-voiding contractions, or later parts of a bladder fill cycle can increase bladder capacity significantly, sometimes to the same level as continuous stimulation [17]–[21], [48]. Potts et al. found in rats that SNM only in the second half of the bladder fill cycle increased bladder capacity significantly, while stimulating the first half did not [48]. Wenzel et al. found that pudendal nerve stimulation at the beginning of bladder contractions increased bladder capacity twice as much as continuous stimulation [17]. Similarly, in our study, closed-loop stimulation was dependent upon bladder contractions, and there was more stimulation in the second half of the bladder fill as bladder contractions became more frequent (Figure 4) and this was fundamentally different than an intermittent stimulation paradigm (i.e. stimulation applied in a 10 s on/off schedule) that does not utilize any sensory feedback. Clinical studies of dorsal genital nerve stimulation also suggest that stimulation only after the urge to void, for as short as 30s in duration, can lead to mean subjective improvements of 73% in the incontinence score [21]. Compared to these closed-loop strategies, our method does not require patient intervention and uses a sensor implanted near the stimulation site.

An increased NVC frequency or a decreased NVC interval are often associated with overactive bladder in pre-clinical models [49]. While it is unclear whether NVCs occur more often in human patients with OAB, NVCs can activate the same bladder sensory neurons as voiding contractions [31] and can therefore elicit an unnecessary urge for voiding that needs to be suppressed. In this study, we did not find any significant impact of closed-loop stimulation on peak pressure, number of non-voiding contractions, or non-voiding contraction intervals. This lack of statistical significance could be a result of low sample size; however, we did not expect the peak pressure to increase as a result of stimulation, and our study results were consistent with this expectation. We hypothesize that this outcome is because SNM relaxed the detrusor muscle, rather than tightened the sphincter muscle, which will lead to higher peak pressure. As a result, the overall bladder volume increased while peak pressure stayed consistent across trials.

Five Hz stimulation was chosen for SNM based on frequency optimization trials that were performed in a pilot experiment. The selection of this frequency is consistent with prior studies which showed that 5 Hz dorsal root stimulation was optimal for increasing bladder capacity when compared to other frequencies [35], [37], and similar to results from another study that concluded 7.5Hz or 10Hz are optimal in minimizing iso-volumetric contractions [45]. Our use of a stimulation amplitude above motor threshold is common in anesthetized experiments (e.g. [45] achieved similar outcome measures as here; others use twice motor threshold or higher in animal studies [37], [38]), as it helps mitigate against the suppressive effects of anesthesia. A previous pudendal nerve stimulation study in behaving felines had greater bladder outcomes at a stimulation amplitude that was twice motor threshold, which was well tolerated by the animals when the amplitude was ramped up [50]. Similarly, ramping up and down of SNM stimulation amplitudes has been described as mitigating uncomfortable sensations of stimulation initiation and cessation in clinical subjects [22]. Brink et al. [51] reported that supra-motor threshold SNM stimulation that increased bladder capacity was successfully used and tolerated in fully conscious large animals. Additionally, this study used healthy animals, which may have contributed to the need for higher stimulation amplitudes as compared to an OAB or simulated OAB model.

The NRMSE and R decoding performance (Table 1) for no-stimulation trials was an improvement upon (NRMSE) or consistent with (R) our previously bladder pressure decoding study (0.28 ± 0.13 and 0.84 ± 0.19, respectively) [31]. As anticipated, closed-loop stimulation increased the NRMSE (0.29, similar to [31]) and decreased the R for bladder pressure estimation, as additional threshold crossings were detected during stimulation (due to possible stimulation artifacts and/or stimulation driven units). Refinement of our cross-channel artifact invalidation may be necessary. Additionally, stimulation itself may have led to a reduction in bladder sensory neuron sensitivity (Figure 6), which would have decreased decoding efficacy during SNM trials. However, it is unclear if sensory feedback is critically necessary during stimulation itself and may not have significantly altered the decision-making process of the closed-loop algorithm.

We did not observe a strong correlation between bladder capacity and the NRMSE for pressure estimation (Figure 5a). Stimulation was only triggered by the closed-loop algorithm based on a relative increase in the bladder pressure, therefore the system could tolerate a small prediction error as well as any amount of baseline offset due to a shift in the noise floor. A large absolute error (or a large NRMSE) might not lead to a high error rate in our closed-loop system, but a low correlation coefficient may indicate a possible loss of channels and lead to inaccurate sensory feedback, less efficient stimulation, and ultimately, lower bladder capacity. The positive, non-significant trend between R and normalized bladder capacity suggests that an increase in bladder capacity may be associated with accurate sensory feedback and the timing of stimulation being applied in closed-loop neuromodulation. The accuracy of identifying bladder contractions was higher in the second half of trials, when stimulation was triggered more often (Figure 4) as the NVC rate increased.

A potential hypothesis for a mechanism of action of SNM is that SNM stimulates sensory pathways, bringing down the level of urgency by inhibiting sensory neural firing [37], [52]. A study in cats showed that stimulating sacral-level dorsal roots inhibits isovolumetric bladder contractions, while stimulating the ventral root did not [37], which is consistent with this hypothesis. In our study, we analyzed the effect of SNM on some bladder sensory neurons (Figure 6, Table 2). For these neurons, we found that the pressure threshold increased and the firing rate as a function of pressure decreased during SNM. This could be a result of a direct effect of the stimulation on the bladder sensory neurons (e.g, collision block) or an indirect effect. We think an indirect effect is more likely given that for 7 of the 8 identified neurons the stimulation lead was on the opposite side of the recording array. There are several possible mechanisms including modulation of interneuron firing and inhibition of the preganglionic efferent neurons of the micturition reflex [52]. This was a small sample size and one neuron did not follow the trend of the other units, however we believe this warrants additional follow up and may yield important insights into the mechanism of SNM.

Both pre-clinical and clinical neurostimulation evidence suggests that continuous stimulation may not be necessary to deliver optimal improvement in bladder capacity and incontinence [17]–[26]. Therefore, it is important to minimize the overall amount of stimulation delivered, as long-term chronic stimulation can facilitate neural habituation [15], [16] that potentially reduces SNM effectiveness, or may result in bothersome stimulation [13]. The methods developed in this study are translatable to clinical use. In this study, DRG were accessed with an invasive laminectomy that is unlikely to be performed clinically. DRG can also be accessed percutaneously at the lumbosacral level, as is done clinically for dorsal root stimulation for pain [47]. A similar approach may allow for less-invasive delivery of a non-penetrating or minimally-penetrating microelectrode within the limited vertebral space around a target DRG. Recent research has demonstrated that DRG cell bodies are more likely to be located near the surface of feline and human DRG [53], [54], and that a thin-film DRG-surface electrode can record neural activity from the bladder in felines [55]. Electrodes with a lower profile may be more feasible to implement clinically than those used in this study.

An aim of this study was to increase bladder capacity, as low bladder capacity is one of the primary symptoms in overactive bladder. The anesthetized non-OAB animal model described in this paper has several limitations, including preventing a full evaluation of clinical OAB parameters such as urinary frequency, urinary urgency, incontinence episodes, and other symptoms. The non-dysfunctional bladder state of these animals may have limited the improvements that were possible for the bladder measures. It is also possible that anesthesia had a suppressive effect, or the relatively short intervals between bladder fills with different stimulation types had a carry-over effect. Awake testing with a dysfunctional bladder model across multi-week intervals may eliminate these potential confounds and would enable longitudinal comparisons between continuous stimulation and closed-loop stimulation. Our previous pudendal nerve stimulation study [50] demonstrated the feasibility of performing bladder neuromodulation with stimulation at or above motor threshold while recording urodynamic parameters (e.g. cystometry curve, bladder capacity, and voiding efficiency) in a freely behaving feline model, and in a separate study we have observed bladder units in chronic feline experiments across multiple weeks [56]. Moving forward, experiments using awake, behaving animals may be most useful for evaluating both the acute and chronic effects of closed-loop SNM without the influence of anesthesia, and would allow for assessment of stimulation amplitudes at and below motor threshold.

## 5. Conclusion

We have demonstrated that closed-loop SNM using DRG signals as sensory feedback can lead to a significant increase in bladder capacity in an anesthetized feline model. Our closed-loop approach matched the effectiveness of continuous SNM while using significantly less stimulation time. Additionally, our neural recordings from bladder sensory afferents suggested that SNM may cause a shift in the relationship between bladder sensory neuron firing rates and bladder pressure, which is consistent with a hypothesis that SNM works by reducing bladder afferent activity. Long-term studies with behavioral animal models will mitigate the effects of anesthesia and repeated bladder fills in a short time frame, and will be critical as a bridging translational step prior to clinical studies.

## 6. Acknowledgements

The authors thank Dr. Sarah Offutt and Dr. Thaddeus Brink for providing comments and discussion, and Eric Kennedy, Dr. Ahmad Jiman, David Ratze, Dr. Jessica Xu, Zuha Yousuf, Alec Socha, Vlad Marcu, Manorama Kadwani, Nicholas Peck-Dimit, and Hannah Parrish for assisting with data collection. The authors also thank the University of Michigan Unit for Laboratory Animal Medicine for assistance with animal care. This study was supported by a research grant from Medtronic and awards from the National Institutes of Health SPARC program (OT2OD023873, OT2OD024907). The opinions expressed in this article are the authors’ own and do not reflect the view of Medtronic or the National Institutes of Health. Drs. Zirpel and Bittner are employees of Medtronic.

## 7. Data Availability

Raw data and analysis software code are available at https://osf.io/jq5hn/.

